# A Classification-Based Generative Approach to Selective Targeting of Global Slow Oscillations during Sleep

**DOI:** 10.1101/2023.10.16.562622

**Authors:** Mahmoud Alipour, SangCheol Seok, Sara C. Mednick, Paola Malerba

## Abstract

**Background:** Given sleep’s crucial role in health and cognition, numerous sleep-based brain interventions are being developed, aiming to enhance cognitive function, particularly memory consolidation, by improving sleep. Research has shown that Transcranial Alternating Current Stimulation (tACS) during sleep can enhance memory performance, especially when used in a closed-loop (cl-tACS) mode that coordinates with sleep slow oscillations (SOs, 0.5-1.5Hz). However, sleep tACS research is characterized by mixed results across individuals, which are often attributed to individual variability.

**Objective/Hypothesis:** This study targets a specific type of SOs, widespread on the electrode manifold in a short delay (“global SOs”), due to their close relationship with long-term memory consolidation. We propose a model-based approach to optimize cl-tACS paradigms, targeting global SOs not only by considering their temporal properties but also their spatial profile.

**Methods:** We introduce selective targeting of global SOs using a classification-based approach. We first estimate the current elicited by various stimulation paradigms, and optimize parameters to match currents found in natural sleep during a global SO. Then, we employ an ensemble classifier trained on sleep data to identify effective paradigms. Finally, the best stimulation protocol is determined based on classification performance.

**Results:** Our study introduces a model-driven cl-tACS approach that specifically targets global SOs, with the potential to extend to other brain dynamics. This method establishes a connection between brain dynamics and stimulation optimization.

**Conclusion:** Our research presents a novel approach to optimize cl-tACS during sleep, with a focus on targeting global SOs. This approach holds promise for improving cl-tACS not only for global SOs but also for other physiological events, benefiting both research and clinical applications in sleep and cognition.

## 1. Introduction

Sleep is crucial for health and brain functions, including mood and cognition [1]. In particular, changes in non-rapid eye movement sleep (NREM) are associated with mild cognitive impairment [2], Alzheimer’s Disease [3] and other neurodegenerative issues (e.g., Parkinson’s Disease [4]). Oscillatory events found in the EEG during NREM sleep have been causally implicated in changes in overnight memory performance. Cortical slow oscillations (SOs, 0.5-1.5 Hz), thalamo-cortical spindles (11-16 Hz) and hippocampal sharp-wave ripples (>100Hz) [5, 6], have all been experimentally connected to consolidation of episodic memory, with performance benefitting from enhancement of the oscillations and their coordination, or disruption of these oscillations leading to loss of performance [7-10]. The current mechanistic understanding of the role these oscillations play in memory changes centers on their time-based coordination, with hierarchical nesting of ripples within spindles, and in turn within SOs, hypothesized to support reactivation of memory traces, and hence mediate the activity-dependent reorganization of synaptic connections that strengthen episodic memory (known as Active Systems Consolidation Theory [11, 12]). This is built on research showing reactivation of behavior-linked neuronal activity enhanced during oscillations [13], as well as correlational and causals studies linking these events with episodic memory outcomes [12, 14, 15]. Beyond consolidation of episodic memory, SOs are also relevant to broad cognitive outcomes, as they contribute to the rescaling of synaptic connections, supporting brain homeostasis [12, 16] and glymphatic system clearance [17]. Given their clear importance for cognitive and health functions, research has sought to interact with SOs via brain stimulation to enhance the functional outcomes of sleep. However, the strides in this approach haven’t consistently yielded success [18-21]. We hypothesize that this is due to a failure to account for the space-time properties of SOs during stimulation. Moreover, the omission of inter-individual differences and the use of heuristically established – but not formally optimized – stimulation parameters have hindered the ability to provide consistent results. Our paper introduces a modeling approach that overcomes the limitations of the current approach by constraining the stimulation protocol to properties derived from space-time profiles of naturally occurring SOs.

Beyond cl-tACS, other techniques are used to interact with SOs to enhance memory, including auditory, magnetic, and electrical approaches. Auditory stimulation involves presenting meaningful cues or non-meaningful sounds [22-25]; targeted-memory reactivation presents cues associated with specific memories [26-30]; and transcranial magnetic stimulation employs magnetic fields non-invasively [31-34]. All these approaches have been shown to enhance SO and spindles, and to boost cognition. A variety of electrical approaches are also available, with transcranial electrical stimulation (tES) applying low-intensity electrical currents [37-39], and, depending on the waveform, being categorized as transcranial direct current stimulation (tDCS) [35], tACS [36], slow oscillatory tDCS (so-tDCS) [6], or transcranial random noise stimulation [37]. All these different types of tES improve learning outcomes [38, 39] and influence memory consolidation by increasing the likelihood of action potentials in prefrontal regions, where SOs are generated [40], thereby enhancing SOs and spindle activity [41]. Despite the great potential and ongoing progress, several factors contribute to SO-focused brain stimulation techniques giving rise to highly varying outcomes, and challenges in achieving consistent results [18-21]. InterLindividual differences among participants in anatomical (e.g. skull thickness, cortical morphology) and physiological (e.g. neurotransmitter availability and receptors distribution) traits can impact the response to tDCS, and tACS [42, 43]. Moreover, for any stimulation technique, the optimal timing of stimulation and number of stimulation sessions are still subjects of ongoing research and debate [44-46]. Determining the ideal stimulation parameters, including current density, duration, and waveform, remains a challenge [47, 48]. Additionally, the heterogeneity in experimental design across studies (e.g., sample size, age and sex of participants) can contribute to difficulty in consolidating experimental outcomes in meta-analytical studies [49]. Another potential confound is introduced when stimulation techniques targeting SO dynamics do not explicitly consider the space-time properties of the SOs. We argue that to improve the consistency in stimulation outcomes, protocols need to consider the variability in individual response to stimulation and the space-time dynamics of SOs during stimulation. In this study, we present a modeling approach for SO-based cl-tACS built on the space-time properties of the SOs during stimulation, which can be leveraged for personalized tailoring of stimulation protocols. We believe that this approach has the potential to address these issues effectively.

The cl-tACS protocol we focus on targets a specific space-time profile of SOs: global SOs. In recent work [50], we showed that a data-driven approach could categorize SOs on the scalp as global, local, or frontal, based on their presentation across electrodes. Global SOs, observed across nearly all electrodes with short delay, exhibited higher trough amplitudes compared to other SO types and demonstrated stronger coordination with sleep spindles. In a follow-up study [15], we reported that global SOs facilitated information transfer over long distances, as evidenced by the strong correlation between effective connectivity estimates and episodic memory improvement across a night. Other, non-global, SO types did not exhibit these characteristics or show significant relationships with memory. These findings underscore the importance of considering the space-time dynamics of SOs when aiming to enhance memory through brain stimulation techniques, and provide the rationale for the current study to focus on establishing a stimulation paradigm capable of selectively enhancing global SOs.

A tACS protocol consists of the electrode montage, waveform, and duration of the stimulation. To achieve better spatiotemporal resolution, we divided the duration of the stimulation into smaller time windows (Δt). This provided more precise control over the timing of targeting specific brain regions. To shape a stimulation protocol to target global SOs, we first generated a set of stimulation paradigms and used forward modeling to simulate brain dynamics during those paradigms. We optimized the parameters for each paradigm to match the profile of an average global SO extracted from sleep EEG. Finally, a classification-based approach was used to select the stimulation protocol among all paradigms considered. This study provides a framework for developing personalized brain stimulation protocols based on individual specific space-time profiles of SOs.

## 2. Materials and Methods

Our model of a global SO targeting protocol is achieved through two procedures: data processing and simulation. Data processing (Fig 1A) of sleep EEG includes generating the average global-SO current density (CD) representation as a time-by-region matrix, and training a classifier to label the CD profiles of SOs as global or non-global. The simulation stage (Fig. 1B) includes defining a search region for stimulation protocols (electrode montage, waveform, and Δ*t*) and identifying the optimal protocol within this region. Parameter tuning for each waveform relies on the average global SO CDs and the global/non-global SO classifier for labeling stimulation paradigms.

**Figure 1.**
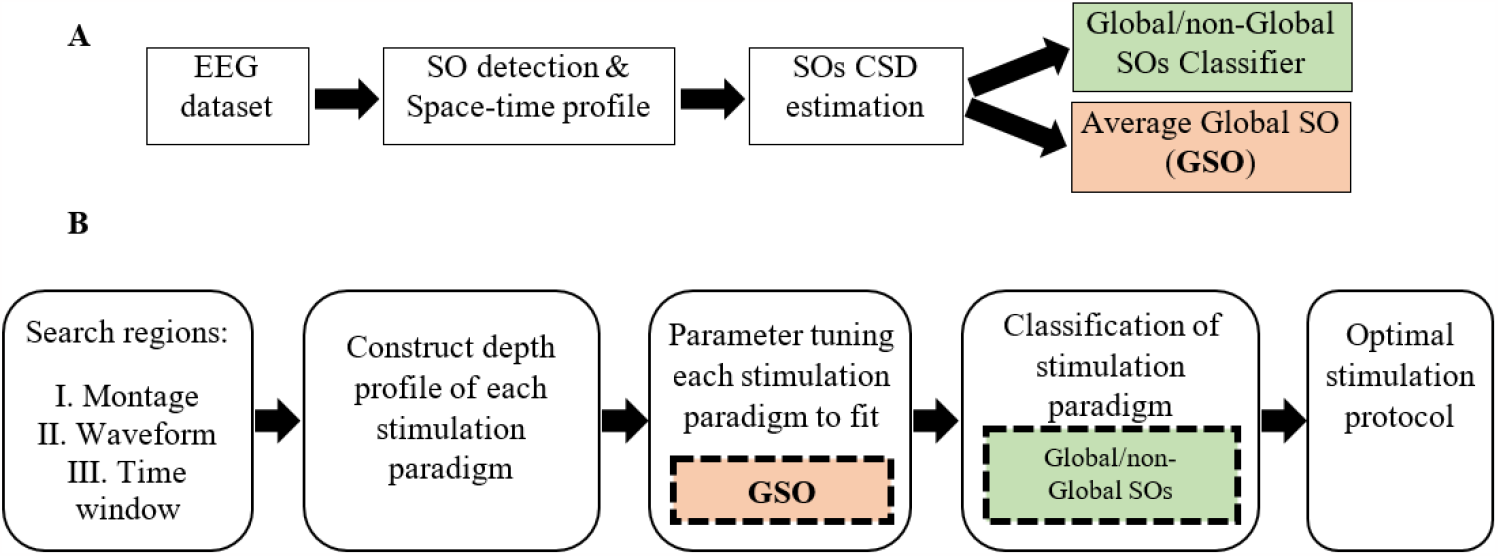
The Global-SO-targeting protocol. A) Sleep data processing procedure, producing a global/non-global SOs classifier and the average CD of global SOs. B) Simulation of stimulation procedure. The parameter tuning stage leverages the average global SOs CD. Classification is based on the global/non-global SOs classifier trained on sleep data.

### 2.1. The EEG dataset

The present study is based on a dataset of sleep polysomnography from the Sleep and Cognition Lab at University of California Irvine, led by Dr. Mednick. The dataset is introduced in detail in [51]. The dataset includes full-night EEG sleep data from 22 healthy volunteers (9 females) without psychological or neurological issues. EEG signals were recorded using a 64-channel cap based on the international 10-20 System at a 1,000 Hz sampling rate, which was subsequently down-sampled to 128 Hz. Out of the 64 channels, 58 recorded the head signal, while others served as reference, ground, and other biosignal channels. Sleep stages (Wake, Stage 1, Stage 2, SWS, and REM sleep) were visually scored in 30-second epochs following the R&K manual [52], using the MATLAB toolbox HUME [53]. Participants all had a good quality sleep (basic sleep characteristics in Supplementary Table S1).

### 2.2. SO detection and space-time profiles

To detect SO events, we used an algorithm previously used in [51] that closely followed the criteria introduced by Massimini et al. [54] and Dang-Vu et al.[55]. SO detection was performed at each electrode independently, discarding times where an artifact was found. The algorithm relied on well-established conditions regarding the amplitude and duration of SO events, and a comprehensive description can be found in prior publications [50, 51]. After SO detection, we performed clustering of SO co-detections to reveal the space-time patterns of SOs, with the same method used in [50, 51]. Briefly, we built separate co-detection matrices for S2 and SWS, where each SO at each electrode was used to generate a binary array the length of all available head electrodes, allowing for a delay of 400ms for co-detections. The co-detection binary matrix was then analyzed with k-means clustering (k=3) using Hamming distance, with 200 replicates and a maximum iteration of 10,000. The clusters were labeled Global, Local or Frontal based on the representation of their centroids on the scalp. Counts of detected SOs in S2 and SWS per each participant, and of global and non-global SOs, are reported in supporting Table S2.

### 2.3. SO current source density estimation

Our study required an estimate of current source density (CD) for each detected SO, which we performed analogously to our previous work [51]. To achieve this step, data from all EEG channels from 500ms before to 500ms after the trough of the SO were binned at Δ*t* time scale and then imported into Brainstorm [56]. Data import to Brainstorm was done for Δ*t* values of 20ms, 50ms, 100ms or 200ms, separately. Since in this retrospective dataset we did not have MRI for the participants, a mixed head model was created with both cortex and sub-cortical substructures using the MNI ICBM152 package, which is highly compatible with different properties within Brainstorm. Because this study focuses on SO dynamics, we included in the model regions that are known to be involved in SOs directly and regions that are potentially involved in coordinated network activation during SO dynamics. The regions included in the model were: neocortex, hippocampus, nucleus accumbens, amygdala, brainstem, caudate nucleus, putamen, pallidum, and thalamus. All regions but the brainstem were considered separately in their left and right hemisphere components. Source estimation was performed at each time point of interest by fitting current dipoles in a fixed three-dimensional grid composed of voxels with 15,002 vertices for neocortex and 5,095 vertices for sub-cortical structures. The boundary element method (BEM) OpenMEEG was used to compute the lead field matrix [57], and the minimum norm method was used to estimate a solution to the linear inverse problem. The identity was used as the noise covariance matrix, and standardized low-resolution brain electromagnetic tomography (sLORETA) was applied, due to our interest in subcortical regions activation, to obtain a final current source density value in each voxel [58]. This allowed for comparison of CD profiles across individuals and minimized bias in the source estimates. In this representation, the CD of each SO is encoded in an m-by-n matrix, where m is the count of Δ*t* bins in a one second time window around the through of the SO (e.g., for Δ*t =* 20*ms*,m is 50) and n is the number of selected brain regions (17 in our case). Thus, the matrix representing spatiotemporal information of CD changes from 500ms before to 500ms after the trough of one SO at Δ*t =* 20*ms* has 850 elements.

### 2.4. SO classification

We classify Global/non-Global CD of SOs with the Bagged Trees algorithm, an ensemble classifier that uses bootstrap aggregation, thus reducing overfitting and improving generalization. Our classifier was built with a bagged ensemble of 100 regression trees. Bagged Trees estimates the probability of an instance belonging to each class by averaging the probability estimates of each individual decision tree in the ensemble. The predicted class is the one yielding the largest probability average. We used 5-fold cross validation for performance evaluation, and 5-fold cross validation in the training datasets. Bayesian optimization was used to find hyper parameters. Features of the classification were the CD of each brain regions in each specific time bin of the CD (i.e., each entry of the CD matrix). We trained two separate classifiers for SOs in stage 2 and SWS. Then, the output of testing each trained classifier on the dataset was organized in a confusion matrix, comprising ‘true positives’ (TP), ‘true negatives’ (TN) and ‘false positives/negatives’ (FP, FN), respectively. Due to the imbalanced nature of the dataset (non-global SOs are more frequent, comprising approximately twice the count of global SOs in our dataset (Table S3)), we use Matthews Correlation Coefficient (MCC), which is highly informative for imbalanced binary classification [59], as our metric for performance. MCC ranges from -1 to 1, with higher values indicating better performance. MCC is calculated using the following formula:

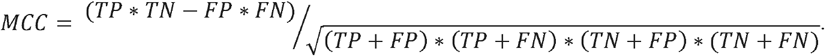

### 2.5. Estimating the current delivered by a stimulation paradigm

In our forward modeling, we estimated the current delivered by a stimulation paradigm, and optimized its parameters to approximate the CD representation of the average global SO found in natural sleep. To estimate the current delivered by any given paradigm, we utilized ROAST, which is a fully automated, Realistic, vOlumetric Approach to Simulate Transcranial electric stimulation [60]. It is an open-source MATLAB toolbox that generates a finite-element model (FEM) mesh and solves the FEM for voltage and electric field distribution in the brain at 1 mm resolution. We used an MNI 152 head to build a transcranial electric stimulation model. In this study, we utilized ROAST to estimate the CD of specific brain regions with respect to the placement of stimulation electrodes, their geometrical dimensions, and initial voltage.

### 2.6. Genetic algorithm

We optimized the parameters of the stimulation paradigm to closely approximate the CD of a global SO. To achieve this, we employed a genetic algorithm (GA): an evolutionary optimization method that iteratively modifies a population of individual solutions [61]. The GA randomly selects individuals from the current population for each generation, with the goal of evolving the population toward an optimal solution. It relies on an objective function to assess how closely a design solution aligns with the desired goal. In our study, the objective function compares the CD of stimulation to the average CD of global SOs in the data, to minimize their difference. In our study, the GA terminated if either the average relative change in the best fitness function value over 50 generations was less than or equal to 1e-6 or if the maximum number of iterations, set at 100 times the count of parameters to be optimized, was reached.

### 2.7. Performance Metrics

The stimulation paradigm that best approximated the CD of an average global SO, ranked based on the posterior probability of classification, was selected as the stimulation protocol. In addition to the posterior probability of classification, which indicates the accuracy of the classification, we employed two additional criteria to evaluate the simulation: the weighted correlation coefficient (WCC) [62] and the weighted mean square error (WMSE) [63]. The WCC measures the similarity between the CD resulting from the stimulation protocol and the data, while the WMSE quantifies the error of the simulation. In the following formula *x, y*, and *w* are data, prediction, and weight (importance), respectively. *n* is the count of features in CD of the data. The WCC is computed as below:

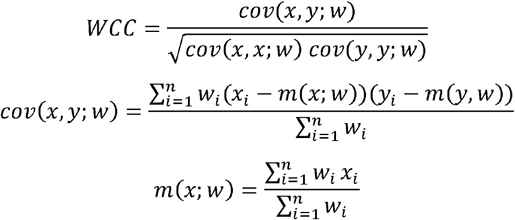

The WMSE is computed as below:

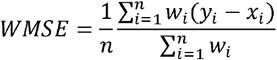

## 3. Results

This study aims to build a principled brain stimulation protocol based on the tACS technique by simulating the currents delivered by tACs and tailoring the protocol parameters to best match the currents naturally found in the sleeping brain during the target event. Simulating a stimulation protocol involves several steps including defining a search set of stimulation paradigms, estimating the currents each paradigm would evoke as function of the parameters, and tuning their parameters to match currents found in the sleeping brain. The last steps include classifying all the stimulation paradigms (either matching the target event of not), and finally finding the optimal stimulation protocol based on classification outcomes.

### 3.1. Stimulation paradigm

To determine the optimal stimulation paradigm, we explored various parameters of a stimulation protocol. Thus, we generated a set of stimulation paradigms by specifying search regions for the electrode montage, stimulation waveforms, and Δ*t* (i.e., the time resolution that we used to sample time in our simulations and data analysis). To limit the search space, we restricted the potential combinations of stimulation paradigm parameters to four electrode sets, four Δ*t* values, and four different waveforms (Fig. 2). For Δ*t*, a particularly relevant parameter that imposes a time scale at which brain dynamics during an SO can be considered stationary enough that it can be represented by its average, we considered 20, 50, 100, and 200ms. For each electrode montage (Fig 2A), we required four electrodes, each chosen from a different quartile of the scalp. We restricted our search region to the electrodes from the 10-20 standard electrodes in each quartile and disregarded electrodes located in the central region. The rationale for selecting one electrode from each quartile to compose any given montage in our search region was to ensure a practical use of stimulation electrodes in real-life implementation of our model. The 10-20 placement preference satisfied two complementary needs: first, placing two electrodes too close to each other could present issues for ROAST when setting up the boundary conditions for the FEM; second, choosing electrodes outside the MRI image boundary could reduce the accuracy of the results. Combining these choices, our search set consisted of 2041 possible montages for active electrodes. For each combination of electrodes, we required two having a positive voltage and two having a negative voltage, such that the sum of all electrode voltages was zero. For the stimulation waveforms (Fig 2B), we allowed for: sinusoidal (W1), sum of three sinusoidal waves (W2), square wave (W3), and a 3^rd^-order polynomial wave (W4). We reasoned that, while not complete, this set of waveforms could capture subtleties in the dynamics of naturally occurring SOs. Each waveform is parameterized with a range of 4 to 10 values, and we aimed to determine the optimal parameters with global optimization approaches. Table 1 illustrates the waveform functions.

**Figure 2.**
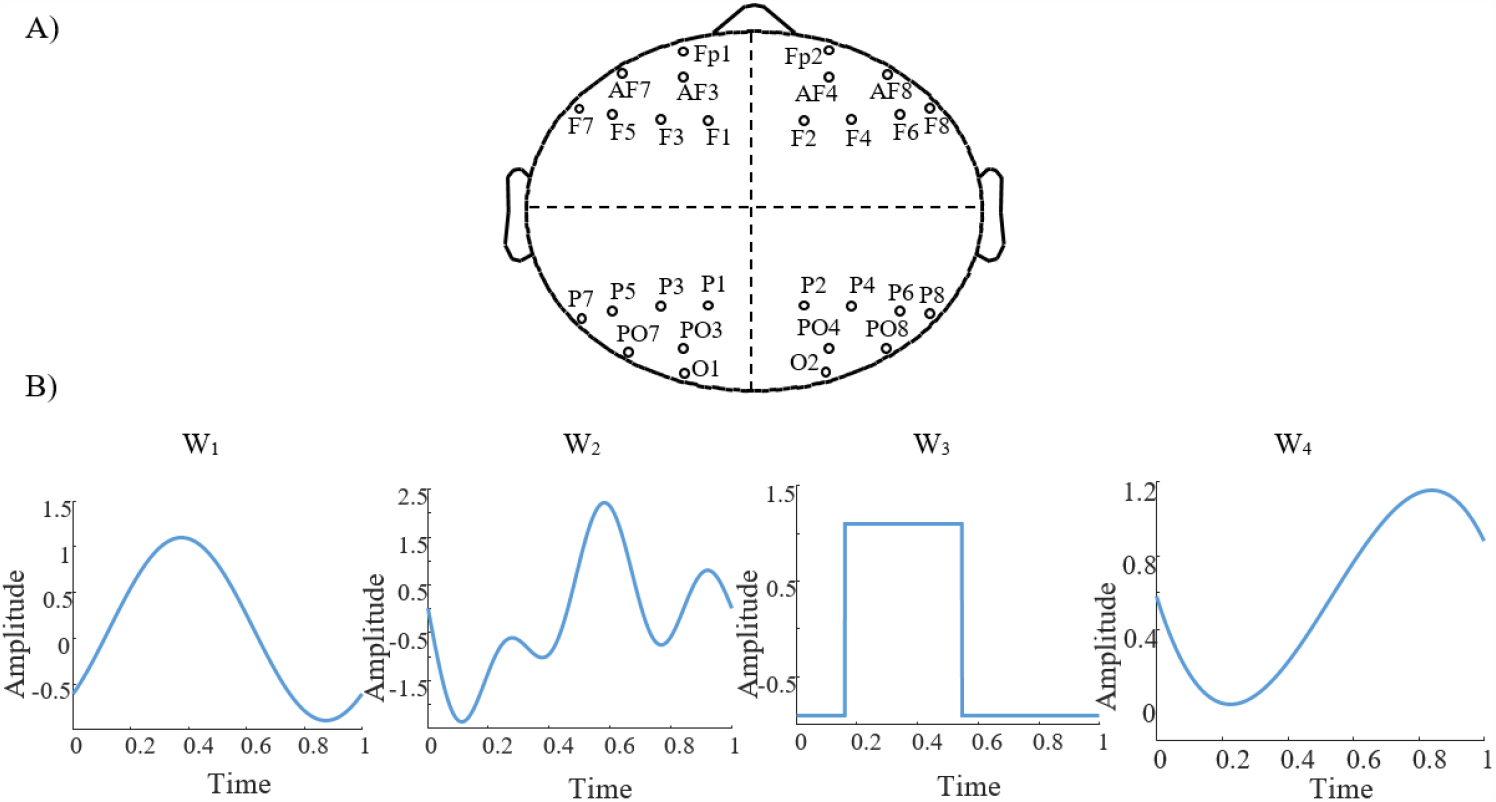
Search region of stimulation paradigm. A) The four regions of choice to search for electrode montage. B) The four stimulation waveforms, W_1_ to W_4_ that constituted our search region for waveforms, each shown here with arbitrary parameters.

**Table 1.** The parameters in the table are amplitude (*A*), frequency (*f*), phase (Ø), offset (*O*), duty cycle (*D*) for square function and coefficients of polynomial function (*P*_1_ to *P*_4_).

| Symbol | formula | Parameters |
| --- | --- | --- |
| $W_1$ | $A \sin(2\pi f t - \phi) + O$ | $A, f, \phi, O$ |
| $W_2$ | $A_1 \sin(2\pi f_1 t - \phi_1) + A_2 \sin(2\pi f_2 t - \phi_2) + A_3 \sin(2\pi f_3 t - \phi_3) + O$ | $A_1, A_2, A_3, f_1, f_2, f_3, \phi_1, \phi_2, \phi_3, O$ |
| $W_3$ | $A \times \text{square}(2\pi f t - \phi, D) + O$ | $A, f, \phi, D, O$ |
| $W_4$ | $p_1(x - \phi)^3 + p_2(x - \phi)^2 + p_3(x - \phi) + p_4$ | $p_1, p_2, p_3, p_4, \phi$ |

### 3.2. Depth profile construction of stimulation paradigm with forward modeling

Any tES protocol, to target a physiological event, delivers definite electrical voltage on the scalp in a short period of time. This process elicits a specific current density across brain regions. To target global SOs, we simulate a stimulation protocol and tune its parameters to elicit a CD in the brain that closely matches the CD of an average global SO. We estimate the CD of any given stimulation paradigm with forward modeling, focusing on a selection of brain regions and time range from 500ms before to 500ms after the SO trough.

Firstly, to numerically encode the stimulation waveform, we chose one waveform from Table 1 and defined it within the range of [0, 1] second, encoded at very high time resolution (0.1ms). The choice of a 1s-long duration for our stimulation protocol was driven by knowledge of the application of the stimulation protocol, in our case to match global SO dynamics both as spatial and temporal events. Spatially, the propagation of SOs from frontal to occipital regions requires about 360ms [54]; temporally, the median duration of global SOs is about 1.0s [50] (Table S3). Hence, a 1s-long duration for the stimulation waveform would allow for an effective realization of a global SO-like dynamics. We then ‘binned’ the waveform in time bins the length of one of the Δ*t* available in our search domain (20, 50, 100 or 200 ms), calculating the average amplitude of the waveform within each time bin. Next, we applied ROAST to estimate the CD induced by stimulation in the brain regions we chose to focus on. Figure 3 shows one example of computing estimates for the case of a stimulation waveform W_1_ (sinusoid), Δ*t* = 100ms, and electrode montage at placed at (F3, F4, P3, P4) with initial voltages of (+1, +1, -1, -1) V, respectively. The waveform and its time-binned representation are shown in Fig 3A, Fig 3B shows the CD of all 17 brain regions for the whole 1s-long interval, while Fig 3C shows the estimated CD across regions within one specific time bin. This estimate is obtained with the assumption that the CD evoked by a stimulation protocol will change linearly with the amplitude of its waveform.

**Figure 3.**
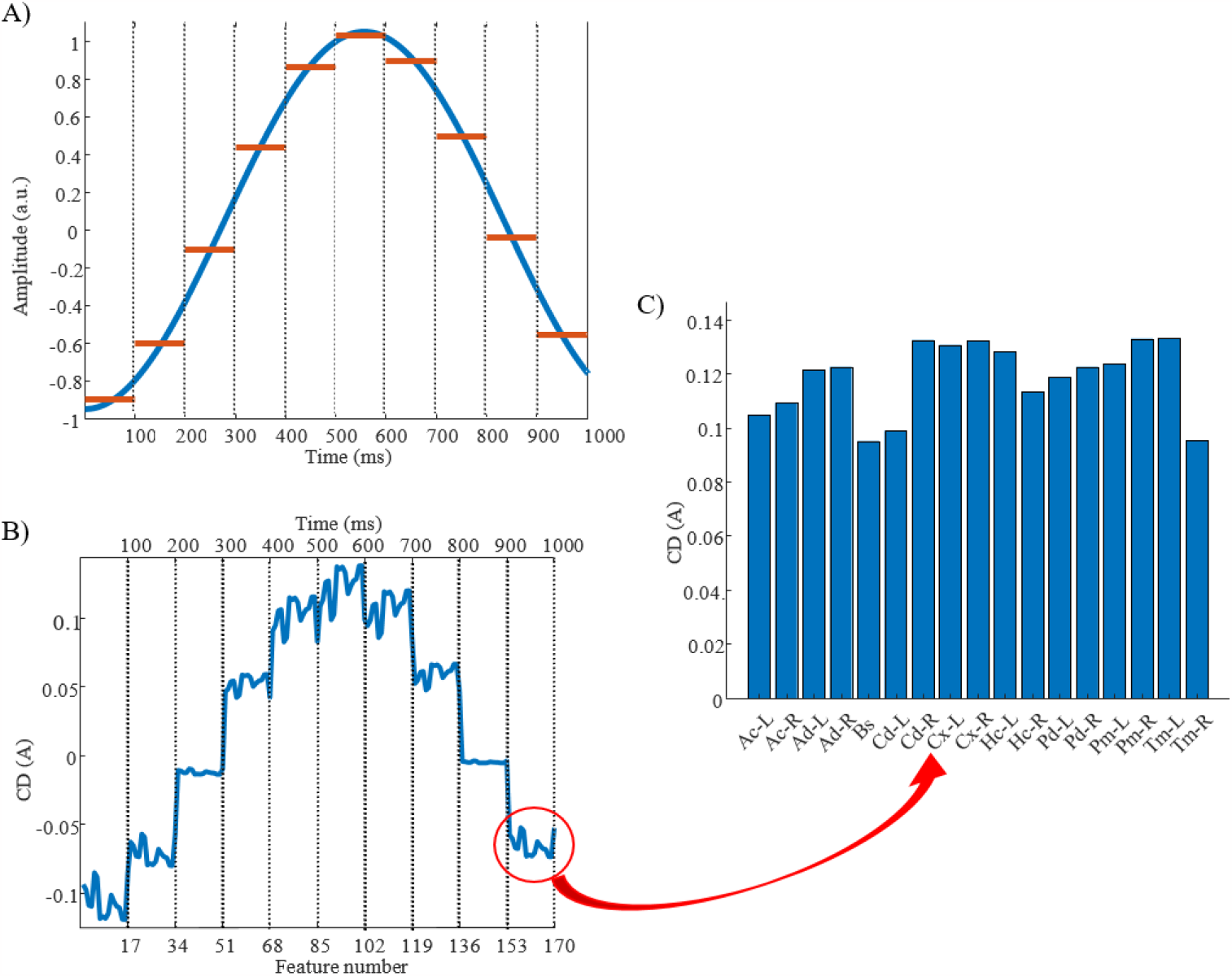
Estimating the CD elicited by a stimulation protocol: example. A) A sinusoidal waveform in the [0, 1] second window with time partitioned in 100ms-long epochs, marked by dashed lines. Blue color shows the waveform with amplitude, frequency, phase and offset equal to 0.2 A, 0.9 Hz, pi/2 s, 0.05 A and red color shows average of the waveform in Δ*t*. B) CD changes of 17 brain regions during one second using F3, F4, P3, P4 as stimulation electrode montage with initial voltages equal to (+1, +1, -1, -1) V, respectively. The x-axis on the top shows time and x-axis on the bottom shows feature number (brain regions) which periodically recur every 100ms proportionally to average of stimulation waveform. The red circle refers to figure C which shows CD of brain regions. C) CD of 17 brain regions. Abbreviations are Ac-L: left nucleus accumbens, Ac-R: right nucleus accumbens, Ad-L: left amygdala, Ad-R: right amygdala, Bs: brainstem, Cd-L: left caudate, Cd-R: right caudate, Cx-L: left cortex, Cx-R: right cortex, Hc-L: left hippocampus, Hc-R: right hippocampus, Pd-L: left pallidum, Pd-R: right pallidum, Pm-L: left putamen, Pm-R: right putamen, Tm-L: left thalamus, Tm-R: right thalamus.

### 3.3. Parameter tuning of stimulation paradigm

We tuned the parameters of the waveform of each stimulation paradigm using a GA. The optimization objective was to obtain CD in specific brain regions that closely resembled the average CD of global SOs. As our previous study suggested some differences could be present in the source currents of SOs in different sleep stages [51], we conducted this optimization approach for global SOs found either in stage 2 sleep (a lighter NREM phase) or in slow wave sleep (SWS, a deeper NREM phase), keeping the two estimates separate. In each case, we defined the objective function of GA as shown in the equation below, where *x*_i_ is the feature *i*^*th*^ in CD estimated from the stimulation, 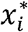 is the *i*^th^ feature in CD of the global SO, *ω*_*i*_ is the weight of *i*^*th*^ feature:

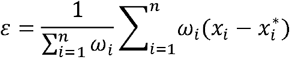

To obtain each feature scaling factor (weight) from the data, we used the F-value of the one-way ANOVA as a univariate feature ranking method [64]. We calculated the F-value between the *i*^Jh^ feature and the global/non-global label of SOs in the dataset. Figure 4 reports the weights of features in Δ t=20ms in SWS, showing that CD in times around the trough are more discriminative of global vs non-global SO. Across all types of waveforms considered, we had to optimize parameters describing amplitudes, frequencies, offsets, and duty cycle (see Table 1). We assigned the parameter search region for the GA as follows: *A* at [0, 1]A for all amplitudes, *f* at [0.01, 4]Hz for all frequencies, Ø at [-π, π for all phases, *O* at [0, 1]A for all offsets, the duty cycle of the square waveform (W_3_) as a percentage value between [0, 100]. Lastly, all coefficients of the polynomial waveform were defined between [-1, 1]. The optimization process finely tuned these stimulation parameters to predict that they would elicit CD in the brain closely replicating the average CD of global SOs. Figure 5 shows the CD of sleep data and optimal stimulation protocol simulation for Δ*t*=20ms, from 500ms before to 500ms after the SOs trough in stage SWS. Figures 5A to 5D show results for waveforms W_1_ to W_4_, respectively. Table 2 shows the desired electrode montage and optimal parameters for each stimulation waveform considered. Figure 5 and Table 2 demonstrate that the CD of EEG and tACS simulation align well, indicating the effectiveness of the chosen electrode montage and waveform parameters in inducing global SOs.

**Figure 4.**
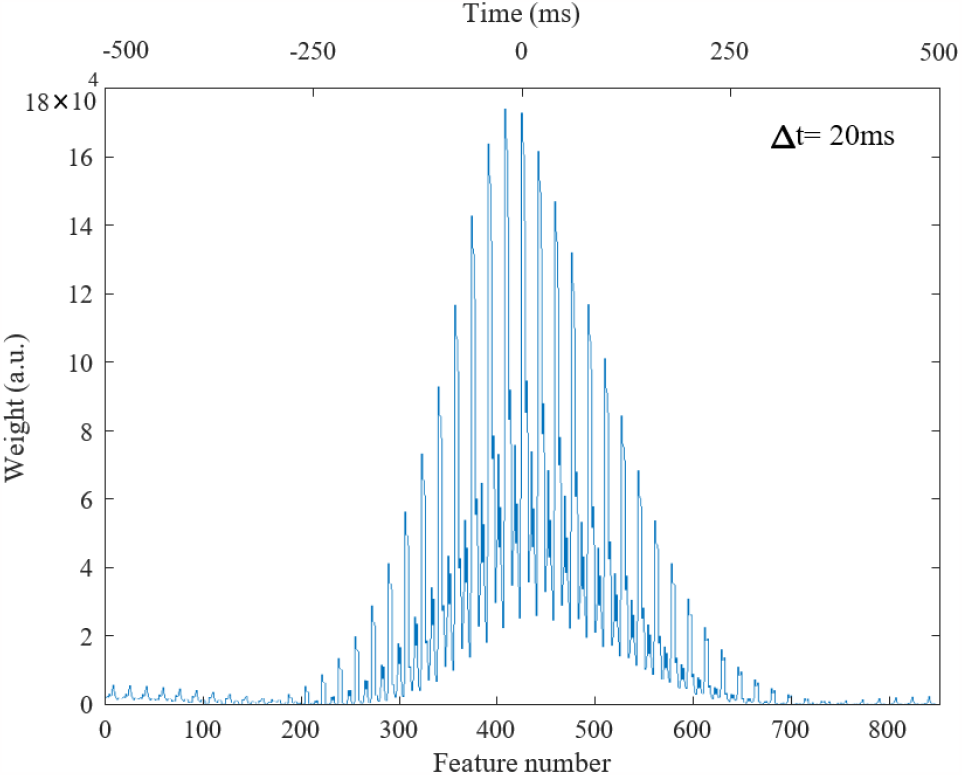
Weights of features in Δ*t*=20ms in stage 3 from 500ms before to 500ms after the SO trough, totally 850 time windows. Lower and upper x-axis are feature number and time (ms), respectively.

**Figure 5.**
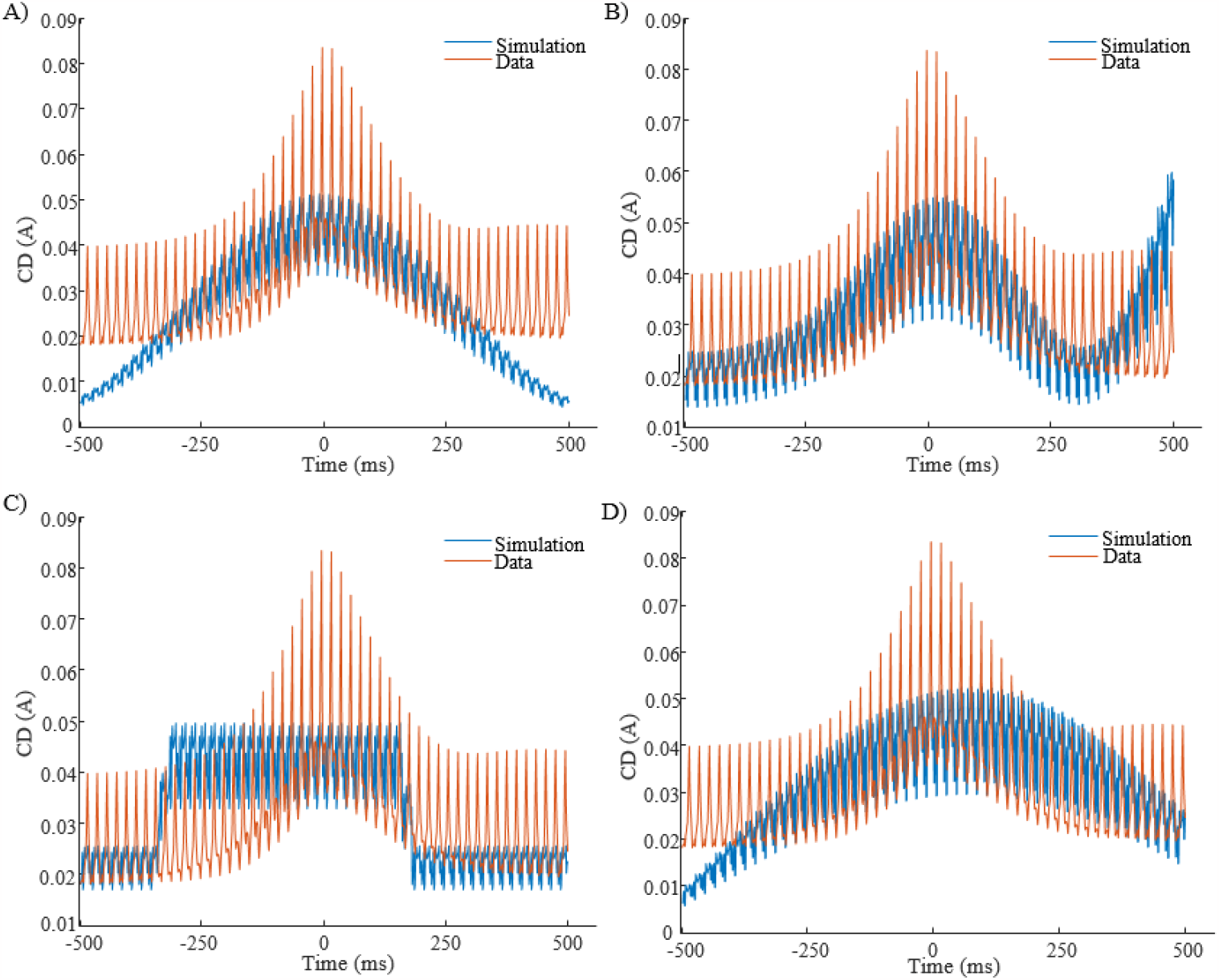
CD of EEG (red color) in stage SWS and tACS simulation (blue color), from 500ms before to 500ms after the SOs trough, for Δ*t*=20ms. Panels are related to waveforms: A: W_1_; B: W_2_; C: W_3_; D: W_4_.

**Table 2.** Optimal electrode montage and waveform parameters for each stimulation waveform in Fig. 5 to target global SOs within the defined search regions.

| Function | parameters | Electrode montage |
| --- | --- | --- |
| $W_1$ | $A = 0.1250$<br>$f = 1.2061$<br>$\phi = 2.3037$<br>$O = 0.2467$ | F7, F2, O1, PO8 |
| $W_2$ | $A_1 = 0.0899$<br>$f_1 = 1.1558$<br>$\phi_1 = -0.6966$<br>$A_2 = 0.1028$<br>$f_2 = 0.1171$<br>$\phi_2 = 1.2957$ | AF7, F6, P2, PO7 |
| | $A_3 = 0.2092$<br>$f_3 = 1.2754$<br>$\emptyset_3 = 2.6255$<br>$O = 0.3500$ | |
| $W_3$ | $A = 0.3825$<br>$C = 0.2441$<br>$\emptyset = 0.5061$<br>$O = 0.0430$ | F7, F4, P1, P4 |
| $W_4$ | $p_1 = 0.5377$<br>$p_2 = 0.6044$<br>$p_3 = 0.5707$<br>$p_4 = -0.2658$<br>$\emptyset = -0.6812$ | Fp1, F6, P3, O2 |

In addition to the optimal electrode montage presented in Table 2, Table S4 provides the optimal symmetric montage, including only symmetric electrodes based on the axis separating brain hemispheres. A choice of symmetric electrode montage can be advantageous in studies aiming to uniformly modulate bilateral brain activity, so we were interested in assessing whether symmetry would provide an advantage. Of note, imposing inter-hemispheric symmetry did not demonstrate a significant difference compared to the optimal generic electrode montage, which can be either symmetric or asymmetric, for each stimulation waveform in Table 2.

### 3.4 Classification of stimulation paradigm

In our modeling approach (Fig 1), once each stimulation paradigm had been parameterized to induce currents as close as possible to global SOs CD, we applied a sleep-trained classifier of global/non-global SO CD to the resulting estimates. The classifier for global/non-global SOs classification trained on the dataset, showed performance measured via MCC at 0.91 in Stage 2 and 0.88 in SWS, respectively. The CD estimate of the current elicited by tuned stimulation paradigm was classified, and, based on the likelihood of belonging to the ‘global SO’ class, the paradigm with the highest posterior probability was chosen as the optimal stimulation protocol. This choice ensures that, among the optimized paradigms capable of targeting global SO, we select the most effective one as the stimulation protocol based on the classifier’s predictions. This approach allows us to identify the optimal protocol by virtue of its close alignment with the global SO characteristics learned by the classifier from EEG data.

To assess the degree of success in matching the current estimated from our optimized protocol with the current underlying the average global SO derived from sleep data, we measured performance metrics including posterior probability of classification, WCC and WMSE (Figure S3). We found that, for SWS, WCC decreases as Δ*t* increases; a trend also seen in stage 2. This was true for all Δ*t* values except at Δ*t*=100ms, where WCC increases. This suggests that our stimulation protocol will yield higher accuracy at smaller Δ*t*. It also indicates that our algorithm could not lead to an effective stimulation protocol for targeting global SOs in stage 2 with square waveform for Δ*t*=100ms.

## 4. Discussion

In the present study we model an electrical brain stimulation protocol to target global SOs using tACS to enhance sleep-dependent memory consolidation. This study introduces a framework of leveraging data about targeted sleep events to determine the stimulation protocol that can best enhance them. This required identifying compatible mathematical representations of current during sleep and current driven by stimulation. Using source modelling, we encoded each SO event in a time-by-region matrix and trained a classifier to distinguish global SOs from non-global ones. Using forward modelling, we estimated the current induced by stimulation paradigms in a time-by-region matrix of the same format as the sleep SO ones. To identify the stimulation current which fitted best the current naturally occurring in the sleeping brain, we built a search domain for all elements of a tACS protocol: electrode montage, stimulation waveform, and sampling time window. We applied a genetic algorithm to optimize these parameters, seeking a combination that closely matched the average global SO current. Finally, by applying our classifier to all stimulation protocols that were optimized, we could choose, among those identified as global SOs, the protocol with the highest posterior probability for classification. We found that the best fit for global SOs in our study is achieved by summing three sinusoidal waves with 10 adjustable parameters and an electrode montage consisting of AF7, F6, P2, and PO7.

The results of this study have several implications. First, we show that targeting global SO can be engineered based on stimulation waveform, duration, and electrode montage. This idea can generalize to the design of stimulation protocols for different applications, beyond our specific interest in sleep global SOs, such as the application of brain stimulation in depression [65] and Parkinson’s disease [66]. Second, our results provide a specific stimulation protocol that can be used in future studies to investigate the effects of global SO stimulation on memory consolidation. Third, our approach can be applied to other SO types (Frontal and Local) to develop targeting stimulation paradigms, which could then be used to causally test their role in cognition in experiments. More broadly, we propose that model-based stimulation paradigms are essential to enable causal evaluation of different space-time presentations of brain rhythms, and can be generalized to other naturally occurring waves, such as sleep spindles.

Including an offset parameter in the waveforms in this study allows the stimulation paradigms, in addition to tACS, to be adaptable for SO-tDCS protocol, which is another method used for memory consolidation [21]. This demonstrates the generalizability of our approach in stimulation protocol design for other types of stimulation methods, beyond tACS. In addition to sinusoidal and square waveforms, which are common brain stimulation signals in literature [67-70], we examine signals of higher complexity, including sum of sinusoidal waveforms and polynomial waveforms, which can be easily generated through Arbitrary Waveform Generators, allowing users to tailor spectral or temporal properties of the signal [71]. Our analysis also showed that the accuracy of the stimulation protocol depended on Δ*t* values, and that there was at least one condition (square waveform, Δ*t*100ms and stage 2 sleep) in which our algorithm could not design a stimulation protocol that targeted global SOs at all, at least within the chosen (and very broad) parameter search regions. In addition to the four different waveforms that we report in this work, we conducted an analysis for targeting global SOs with a Gaussian waveform and found that, while the Gaussian waveform can be used for smallΔ*t*, it is not successful for Δ*t* greater than 50ms. This demonstrates that choosing a waveform requires attention and consideration, and not all waveforms are applicable for brain stimulation. Our approach in stimulation protocol designing, by evaluating the applicability of using a waveform, can help in choosing the desired waveform according to data. This study leverages the assumption that the CD of brain regions can linearly change according to the amplitude of the stimulation waveform, driven by user constraints of ROAST and other finite-element based approaches, such as SimNIBS [72], as they do not consider the initial CD of brain regions that can be calculated from EEG. We also confined our search region for finding the optimal electrode montage to the standard electrodes of the 10-20 system for the sake of problem simplicity. Despite this smaller range, compared to other systems, we still found optimal matches for all waveform types considered. Future studies could increase the number of candidate electrodes to potentially improve the efficiency of the stimulation protocol and possibly its adaptability to a broader range and more complex type of waveforms. This study focused on modeling and optimizing a tACS protocol considering the SO space-time profile of 22 healthy volunteers during sleep. While the present study did not develop personalized brain stimulation protocols, its results provide a framework for developing personalized brain stimulation protocols based on an individual’s specific SOs space-time profile (by isolating the procedures to data only derived from one person). In contrast to a one-size-fits-all approach, personalized brain stimulation protocols are hypothesized to increase the efficacy of the treatment, potentially reducing the duration and intensity of the stimulation sessions, as well as the overall cost of treatment [73-75]. Therefore, studies about developing personalized tACS for memory consolidation should be pursued in the future.

While this study completes the analytical and modeling step of identifying a candidate stimulation paradigm, the efficacy of our stimulation protocol in improving memory consolidation remains to be tested in future experimental studies. Experimentally assessing the similarity/difference between global SO currents and those induced by our global-SO-targeting stimulation is essential. If successful, this study could pave the way to future studies that can validate findings in a clinical setting and investigate the long-term effects of global-SO-targeting stimulation on memory consolidation.

In conclusion, our study introduces a model-based approach to develop stimulation protocols for tES, addressing interindividual variability and targeting specific physiological events in the brain. We achieve this by learning from brain signals and optimizing the simulated current effects of the stimulation. Our results suggest that closed-loop tES can be used not only to target SOs based on their timing and frequency range, but also to consider their space-time profiles. The presented generative approach enables the selective targeting of global SOs, and study provides an optimal stimulation paradigm for future investigations into the effects of global SOs stimulation on memory consolidation. This work, by introducing a novel method for designing stimulation protocols, sheds light on the optimal and efficient approach to brain stimulation.

## Acknowledgements

NIH grant (R01 AG046646) to S.C.M. supported acquisition of data analyzed in this work.

## CRediT authorship contribution statement

### Mahmoud Alipour

Conceptualization, Methodology, Investigation, Visualization, Writing.

### SangCheol Seok

Methodology, Conceptualization. **Sara C. Mednick**: Data acquisition, Conceptualization, Resources. **Paola Malerba**: Supervision, Project administration, Data acquisition, Conceptualization, Methodology, Writing, Resources.

