## Supplementary material for "A Classification-Based Generative Approach to Selective Targeting of Global Slow Oscillations during Sleep"

**Supplementary material**
**Tables**
Table S1. Sleep outcomes for our 22 participants. WASO: wake after sleep onset. For each value, time is reported in minutes.

| | |
|---|---|
| Total Sleep Time | $465.36 \pm 7.76$ |
| S1 | $22.1591 \pm 10.40$ |
| S2 | $199.75 \pm 25.92$ |
| SWS | $126.68 \pm 31.61$ |
| REM | $102.07 \pm 14.96$ |
| Sleep Onset | $14.04 \pm 6.71$ |
| WASO | $5.30 \pm 4.75$ |

Table S2. Count of SOs per participant during stage 2 and SWS.

| | Stage 2 | | | SWS | | |
|---|---|---|---|---|---|---|
| Participant ID | Total SOs | Global SOs | Non-Global SOs | Total SOs | Global SOs | Non-Global SOs |
| 1 | 12016 | 6259 | 5757 | 76568 | 35684 | 40884 |
| 2 | 9263 | 1436 | 7827 | 26372 | 2800 | 23572 |
| 3 | 11244 | 3690 | 7554 | 66459 | 17821 | 48638 |
| 4 | 15677 | 6591 | 9086 | 33090 | 4967 | 28123 |
| 5 | 16835 | 7572 | 9263 | 58246 | 16097 | 42149 |
| 6 | 12962 | 3790 | 9172 | 34982 | 7573 | 27409 |
| 7 | 5649 | 884 | 4765 | 8784 | 1468 | 7316 |
| 8 | 10035 | 4751 | 5284 | 13406 | 4136 | 9270 |
| 9 | 13702 | 4526 | 9176 | 49634 | 17210 | 32424 |
| 10 | 15203 | 7244 | 7959 | 56065 | 20443 | 35622 |
| 11 | 10689 | 2934 | 7755 | 26649 | 7098 | 19551 |
| 12 | 2086 | 0 | 2086 | 4164 | 100 | 4064 |
| 13 | 9911 | 2173 | 7738 | 20128 | 2304 | 17824 |
| 14 | 7224 | 1064 | 6160 | 16009 | 1114 | 14895 |
| 15 | 6917 | 554 | 6363 | 18798 | 2774 | 16024 |
| 16 | 10908 | 6055 | 4853 | 44873 | 17759 | 27114 |
| 17 | 14994 | 8447 | 6547 | 110766 | 58393 | 52373 |
| 18 | 16840 | 8868 | 7972 | 77381 | 32620 | 44761 |
| 19 | 8636 | 1676 | 6960 | 18880 | 3727 | 15153 |
| 20 | 15408 | 7624 | 7784 | 46514 | 16624 | 29890 |
| 21 | 17589 | 9563 | 8026 | 84597 | 42546 | 42051 |
| 22 | 7607 | 1727 | 5880 | 28711 | 8877 | 19834 |
| SUM | 251395 | 97428 | 153967 | 921076 | 322135 | 598941 |

Table S3. Count and median duration (s) of global and non-global SOs during stage 2, SWS, and combined stages.

| | Stage 2 | | SWS | | Stage2 + SWS | |
|---|---|---|---|---|---|---|
| | Count | Median duration (s) | Count | Median duration (s) | Count | Median duration (s) |
| Global SOs | 97428 | 1.1200 | 322135 | 1.0920 | 419563 | 1.0990 |
| Non-global SOs | 153967 | 1.1480 | 598941 | 1.0210 | 752908 | 1.0420 |
| Total SOs | 251395 | 1.2060 | 921076 | 1.0465 | 1172471 | 1.0720 |

Table S4. Optimal symmetric electrode montage and parameters of the stimulation waveforms.

| Function | parameters | Electrode montage |
|---|---|---|
| $W_1$ | $A = 0.1631$<br>$f = 0.0471$<br>$\emptyset = 0.4966$<br>$O = 0.3685$ | F1, F2, P3, P4 |
| $W_2$ | $A_1 = 0.2290$<br>$f_1 = 0.1280$<br>$\emptyset_1 = 1.1215$<br>$A_2 = 0.0801$<br>$f_2 = 1.5466$<br>$\emptyset_2 = -2.4277$<br>$A_3 = 0.0969$<br>$f_3 = 1.1933$<br>$\emptyset_3 = 1.4360$<br>$O = 0.4303$ | F5, F6, P1, P2 |
| $W_3$ | $A = 0.0385$<br>$f = 3.0311$<br>$\emptyset = 1.9479$<br>$D = 52.9285$<br>$O = 0.2919$ | F7, F8, O1, O2 |
| $W_4$ | $p_1 = 0.5377$<br>$p_2 = 0.5707$<br>$p_3 = 0.6044$<br>$p_4 = -0.2658$<br>$\emptyset = -0.6812$ | Fp1, Fp2, P5, P6 |

**Figures**

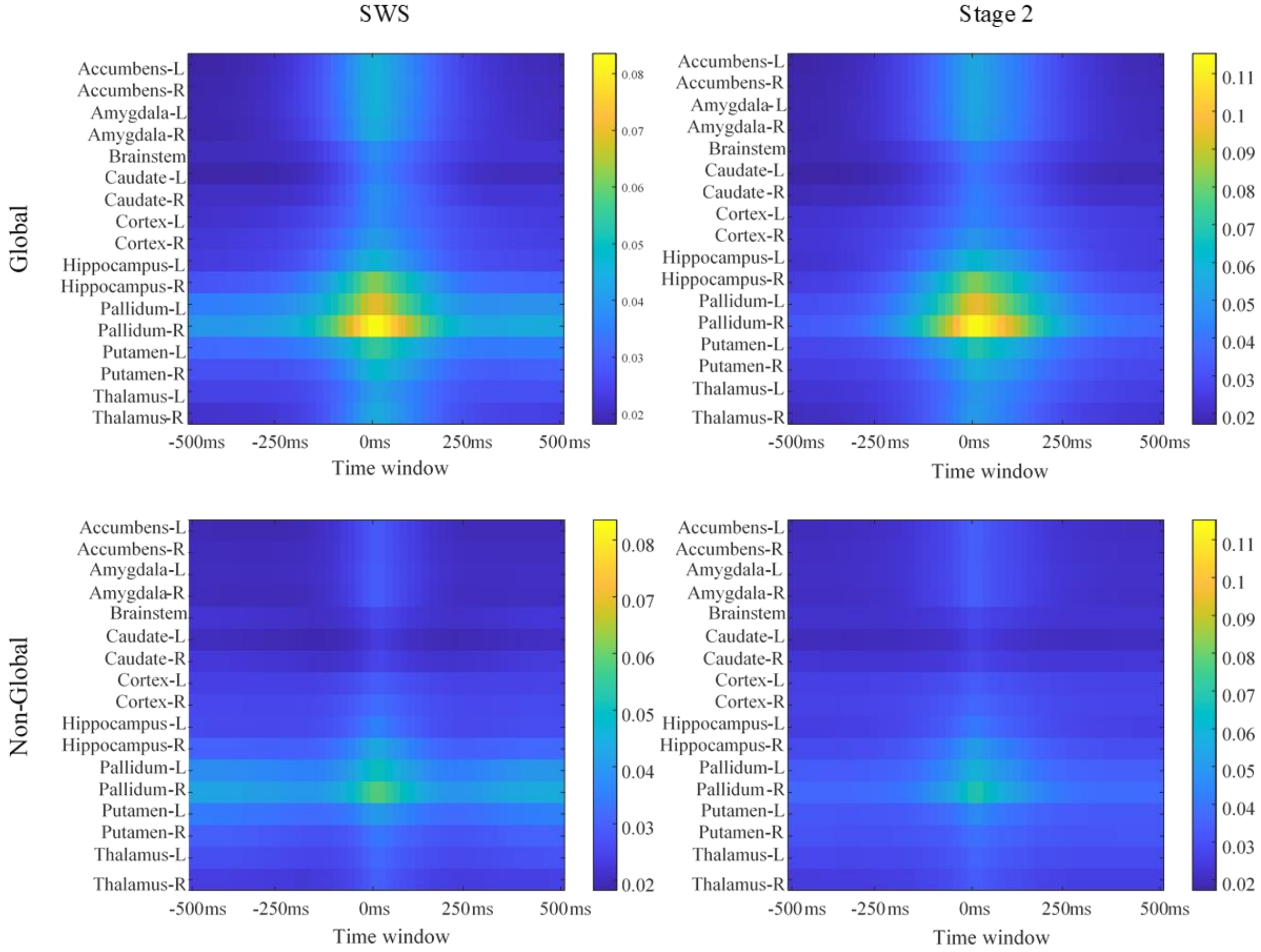


Figure S1. Space-time profile of SOs SWS (left column) and stage 2 (right column) in $\Delta t$=20ms. First and second rows demonstrate space-time profile of global and non-global SOs, respectively.

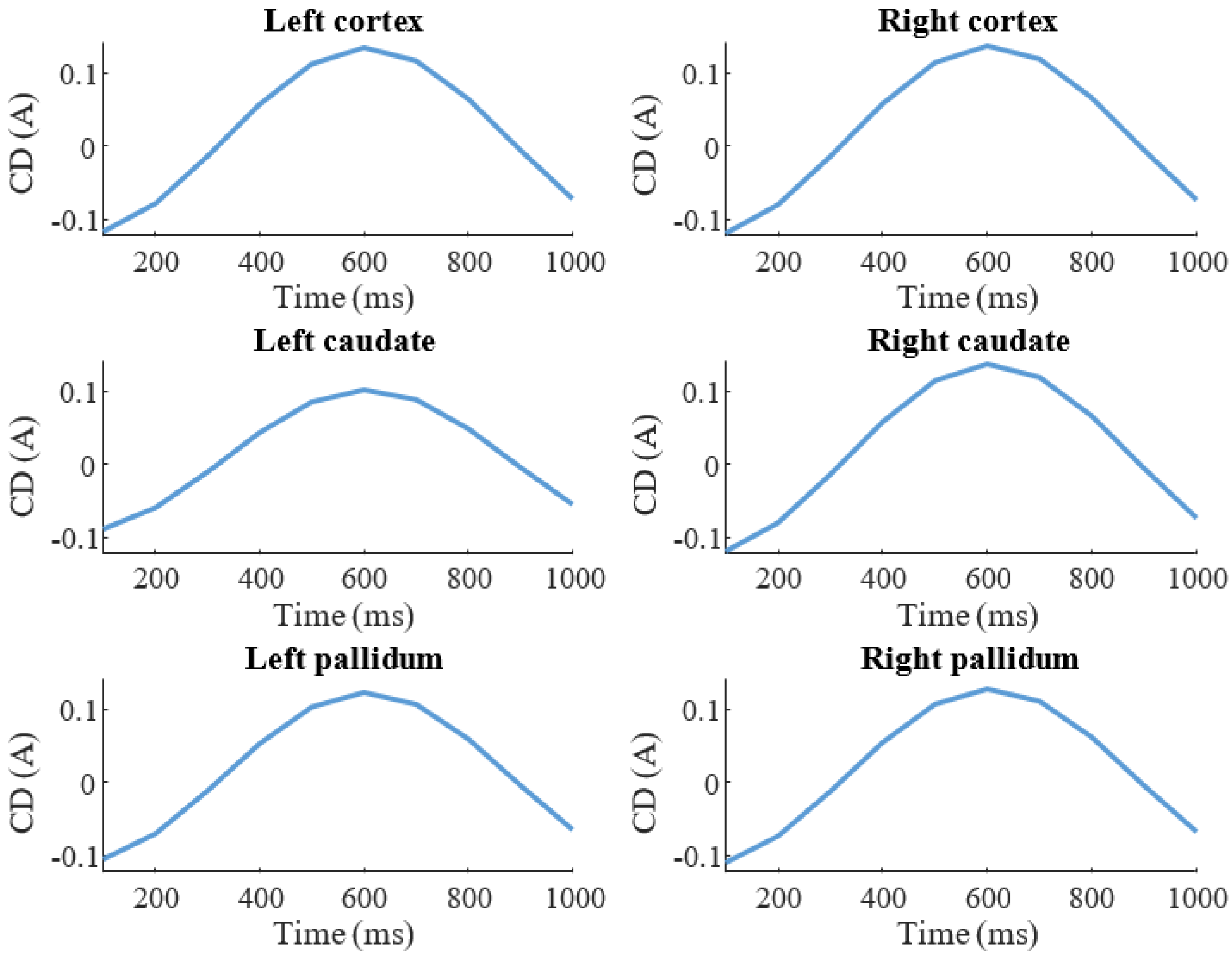


Figure S2. CD of six brain regions during one second of electrical stimulation using $\Delta t$=20ms.

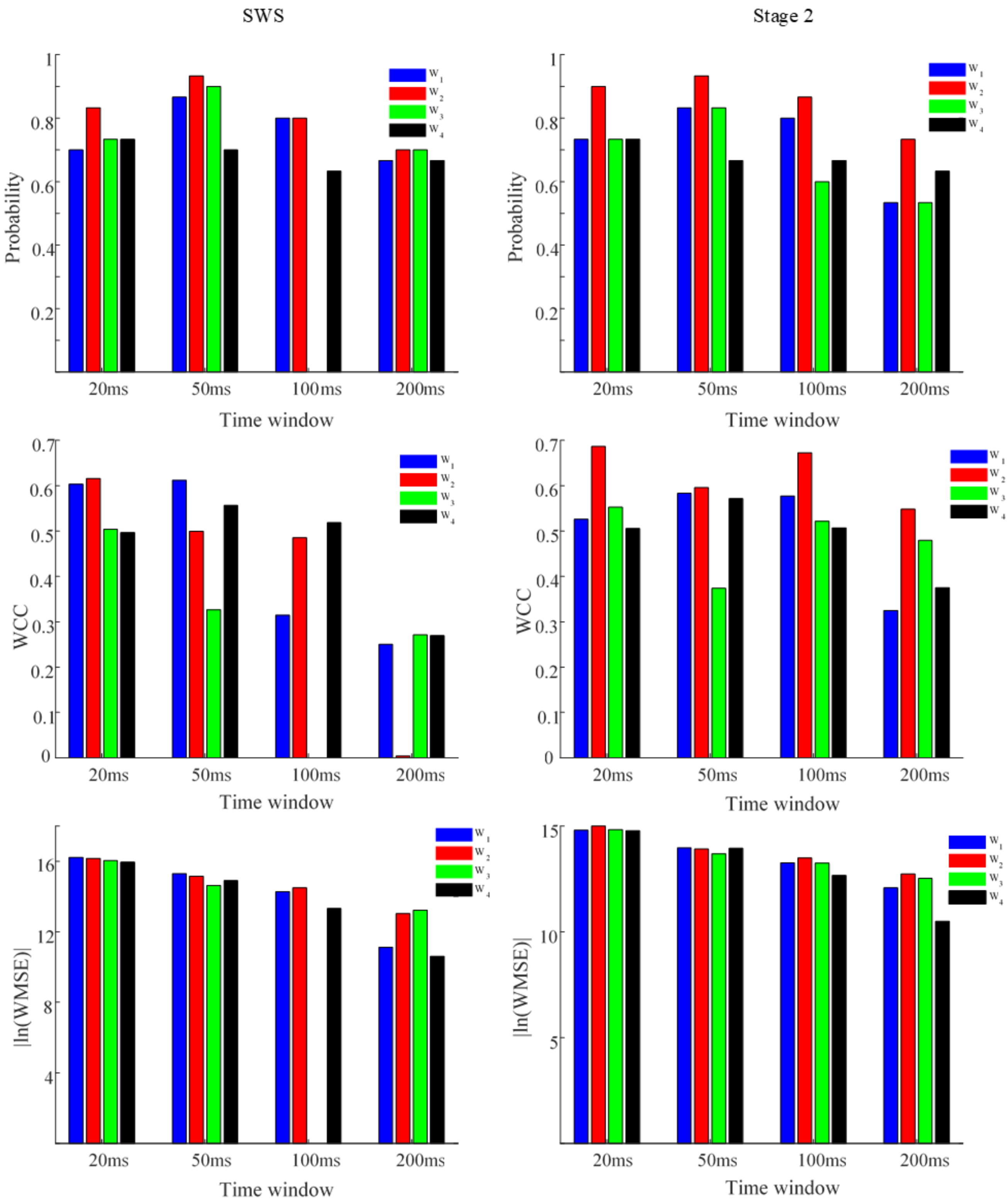


Figure S3. Performance metrics of stage 2 and SWS. First to third rows show probability of classification, WCC and |ln(WMSE)|. Left and right columns show results regarding to stage 2

and SWS respectively. Calculating absolute value of logarithm of WMSE is due to make the values comparable visually.
